# Older adults show overexaggerated and larger noise-related degradation in their neural tracking of speech

**DOI:** 10.64898/2026.07.03.736364

**Authors:** Jessica MacLean, Gavin M. Bidelman

**Affiliations:** Department of Speech, Language and Hearing Sciences, Indiana University, Bloomington, IN, USA; Program in Neuroscience, Indiana University, Bloomington, IN, USA; Cognitive Science Program, Indiana University, Bloomington, IN, USA

**Keywords:** aging, neural synchronization, speech perception, speech-in-noise

## Abstract

**Background:** Speech-in-noise (SIN) perception is a difficult everyday listening task that becomes more difficult with age. Neural tracking of target speech is associated with successful speech perception in clean and noise-degraded listening environments. How aging impacts neural tracking of speech and relates to behavioral decrements in older adults’ SIN perception remains unclear. To address these questions, we measured neural speech tracking during a continuous SIN perception task in younger and older adults via multichannel EEG.

**Method:** Participants (*n*=83) monitored a continuous stream of syllables (~4.5 Hz) presented in quiet and noise conditions during EEG recordings. We assessed neural phase-locking value (PLV) to the acoustic speech envelope to investigate interactions between aging, hearing loss, and stimulus noise on neural synchronization to speech.

**Results:** Compared to younger adults, older adults demonstrated less behavioral sensitivity to noise effects than young adults and had higher overall PLV to target speech. Older adults also showed greater noise-related degradations in neural speech processing relative to younger listeners. Age remained a strong predictor of behavioral responses to speech even after controlling for hearing loss. Covarying for hearing loss removed most age-related effects on neural PLV.

**Conclusion:** Older adults demonstrate overexaggerated neural tracking to ongoing speech presented in quiet and greater noise-related reductions in neurobehavioral speech processing than young adults. Our results support the decline-compensation hypothesis, corroborate unusually large speech envelope encoding in older listeners, and suggest more robust neural synchronization to the speech signal is not always perceptually advantageous.

## 1. Introduction

Everyday listening necessitates the ability to perceive speech-in-noise (SIN), an already-difficult task that becomes even more difficult with age (Heidari et al., 2018). SIN difficulties are a common complaint for older adults (> 65 years of age) even without hearing loss (Cruickshanks et al., 1998; Nash et al., 2011; Pichora-Fuller et al., 2017). Emerging links between declining SIN abilities and risks for social isolation, depression, and dementia highlight the need for interventions which address age-related SIN deficits. However, current behavioral interventions are lacking and hearing aids do not adequately address SIN problems in elderly individuals (Beck et al., 2018). As such, it is imperative to understand the neural and behavioral mechanisms of SIN-related declines in aging to inform the development of targeted interventions.

Difficulties perceiving SIN are perhaps the most common audiologic complaints in the older adult population (Kochkin, 2010). SIN abilities decline exponentially after age 50 due to a variety of age-related changes which impact the auditory system from the cochlea to the cortex, resulting in changes to sensation and perception (Moore et al., 2014). For example, the encoding of temporal fine structure is important for both SIN perception (Hopkins & Moore, 2009) and cortical tracking of speech (Ding et al., 2014), yet decreases with age independent of hearing loss (Hopkins & Moore, 2011). Age-related changes within the central auditory system, including the balance of excitatory/inhibitory activity (Caspary et al., 2008; Caspary et al., 1995) and large-scale neural network connectivity (Martin & Jerger, 2005), could account for SIN deficits in older listeners (Bidelman, Price, et al., 2019; Bidelman et al., 2014).

Though aging and age-related hearing loss (ARHL, presbycusis) both impact SIN processing (Dubno et al., 1984; Moore et al., 2014; Smith et al., 2024), they are difficult to separate. These factors show parallel trajectories of decline and impact SIN processing in similar ways by decreasing access to temporal fine structure cues (Füllgrabe et al., 2015; Hopkins & Moore, 2011) and reducing the perceptual release from fluctuating maskers (Gifford et al., 2007; Gregan et al., 2013). However, age and hearing sensitivity might also be partially separable factors in terms of auditory aging. For example, hearing loss can actually bolster the detection of envelope modulations (Bacon & Gleitman, 1992; Moore & Glasberg, 2001; Wallaert et al., 2017). Yet, enhanced processing of basic amplitude envelope structure does not away transfer to speech, suggesting it may actually distort communication signals and inhibit recognition. Sensorineural hearing loss, independent of age, is associated with decreased peripheral auditory sensitivity alongside central changes in the brain’s white matter (Tarabichi et al., 2018). Such peripheral and central changes in the auditory system could lead to poorer fidelity and internal representations of speech (Bidelman, Price, et al., 2019). From the extant literature, it is clear that both age itself and ARHL reflect overlapping contributions to auditory aging that must be accounted for to fully describe senescent changes in speech and SIN processing.

Neural synchronization, or the phase-locking of neural activity to external stimuli such as speech, is one neural mechanism which underlies successful speech perception (Ding & Simon, 2014; Vanthornhout et al., 2018). Neural synchronization to speech seems to be strongest around 4-5 Hz, the purported “nominal speech rate” across the world’s languages (Assaneo & Poeppel, 2018; Ding et al., 2017; He et al., 2023; Momtaz & Bidelman, 2024). Stronger neural speech tracking relates to improved speech comprehension both in quiet (Riecke et al., 2018; Vanthornhout et al., 2018) and noise (Ding et al., 2014; Etard & Reichenbach, 2019; Keshavarzi et al., 2020). Similarly, deficits in the neural timing of the speech-evoked potentials and unusually large neural tracking of the speech envelope correlate with poorer behavioral speech recognition (Momtaz et al., 2021; Roque et al., 2019; Sidiras et al., 2020), highlighting the importance of temporal encoding for SIN perception.

The ways in which aging and ARHL impact neural envelope tracking and behavioral perception of SIN are not fully understood (Sauvé et al., 2019; Sauvé et al., 2022; Peelle et al., 2010; Criscuolo et al., 2025). Some studies have found that enhanced neural envelope encoding of target speech is beneficial for speech understanding (McClaskey et al., 2019; Petersen et al., 2017; Schmitt et al., 2022). Still, other studies find that enhanced envelope tracking is related to poorer SIN comprehension, perhaps reflecting abnormal “central gain,” decreased inhibition, or increase redundancy of the brain networks subserving speech processing (Bidelman et al., 2014; Goossens et al., 2018; Millman et al., 2017). Differences in target speech material, background noise type, and audibility across studies likely lead to diverging results. However, given the potential for intervention paradigms such as rhythmic cueing or neurostimulation to improve envelope encoding in tandem with speech perception in younger listeners (Erkens et al., 2020; Falk et al., 2017; Guilleminot & Reichenbach, 2022; van Bree et al., 2021; Wilsch et al., 2018) and hearing-impaired adults (Erkens et al., 2021), it is imperative to understand how various factors of the aging process alter the neural encoding of complex speech signals.

In the present study, we investigated age- and noise-related differences in cortical speech tracking in an oddball detection paradigm via high-density EEG. Our dataset comprised a large sample of healthy adults spanning 18-75 years of age and with and without hearing loss (Bidelman, Mahmud, et al., 2019; Bidelman, Price, et al., 2019; Bidelman et al., 2026; Price & Bidelman, 2021). Speech tokens were presented in a repetitive speech stream at the nominal speech rate of 4.5 Hz (Assaneo & Poeppel, 2018) across clean and noise conditions. We measured perceptual sensitivity and neural phase-locking to the running speech to quantify neural-speech synchronization and noise-related changes in speech processing among younger and older adults.

## 2. Methods

### 2.1 Participants

The dataset consisted of N=83 participants aggregated from our previously published EEG studies on speech processing and auditory system function across the lifespan (Bidelman, Mahmud, et al., 2019; Bidelman, Price, et al., 2019; Bidelman et al., 2026; Price & Bidelman, 2021). Participants were split into Younger Adult (YA; *n* = 33, age range: 18-41 years, mean ± SD: 68.6 ± 5.7 years old; 16/17 male/female) and Older Adult (OA; *n* = 50, age range: 52-75, mean ± SD: 23.6 ± 4.2 years old; 15/35 male/female) groups by splitting the participant sample in half at age 50. All participants self-reported no history of cognitive or psychiatric disorders and had no more than moderate hearing loss (pure tone average (PTA) ≤ 45 dB HL) confirmed via audiometric testing (octave frequencies, 250 – 8000 Hz; **Figure 1**). Hearing was symmetric between ears within both groups (*t* test between ears: OA: *p* = 0.1, YA: *p* = 0.09). As expected, older adults had higher PTA thresholds (mean of 500, 1000, and 2000 Hz) than younger adults (PTA_OA_: 21.6 ± 7.9 dB HL; PTA_YA_ : 4.2 ± 4.8 dB HL; *t*(47.7) = 11.3, *p* < 0.0001). None of the participants utilized hearing aids. Cognitive function was not screened. However, we note that mild cognitive impairment actually increases auditory brainstem and cortical evoked potentials to speech (Bidelman et al., 2017) which is not observed in this sample of older listeners (Bidelman, Price, et al., 2019). Other recent studies have also shown that speech envelope tracking does not vary with cognitive status (Bolt & Giroud, 2024).

**Figure 1.**
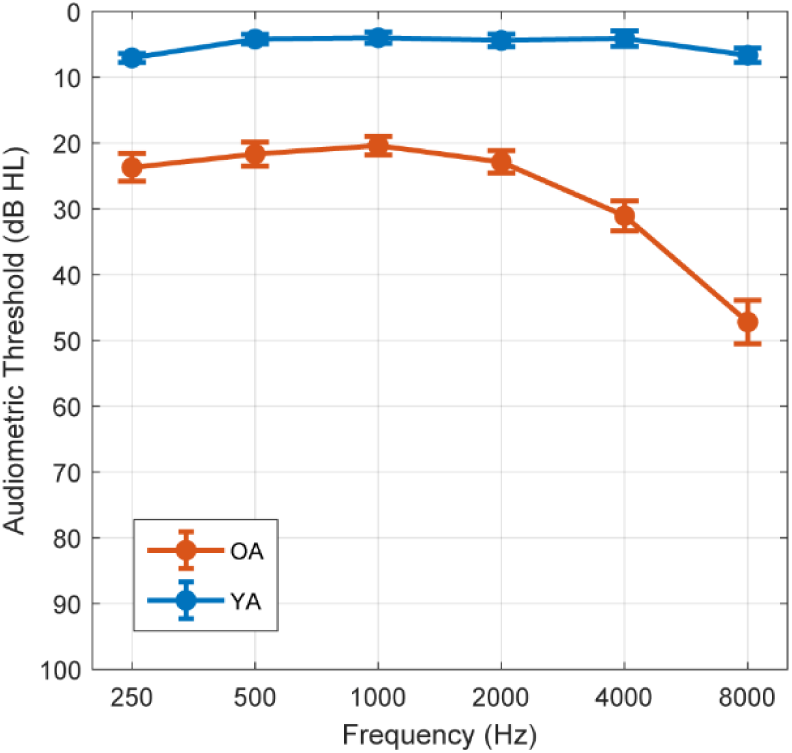
Group-average audiograms. Group average audiometric thresholds collapsed across both ears shown for younger adults (YA) and older adults (OA). OA had higher pure-tone averages (PTA, average of 500, 1000, and 2000 Hz thresholds) than YA (*p* < 0.0001). Error bars: ±1 S.E.M.

### 2.2 Stimuli and task

Specific details of the speech stimuli and task paradigm are described fully elsewhere (Bidelman, Mahmud, et al., 2019; Bidelman, Price, et al., 2019; Bidelman et al., 2026; Price & Bidelman, 2021). Briefly, the primary task consisted of a target speech detection task during simultaneous EEG acquisition (**Figure 2**). Participants detected oddball tokens among a triplet of vowels (/a/, /i/, oddball: /u/ (Price & Bidelman, 2021; Bidelman et al., 2026) or constant-vowels stimuli (/ba/, /pa/, oddball: /ta/ (Bidelman et al., 2019a, b) presented in a continuous stream. Individual tokens were presented in a pseudo-random order so that at least two frequent tokens occurred between successive oddball tokens. There were a total of 2000-3000 standard and 70-210 oddballs, respectively. Each speech token was 100 ms with a fundamental frequency (F0) of 150 Hz. Tokens were presented with a jittered interstimulus interval with an average rate of ~4.5 Hz (i.e., 1/0.225 ms = 4.44 Hz). Speech stimuli were presented in both clean and noise (+5 or 10 dB SNR; 8-talker babble) blocks. In the original studies, SNRs were determined via pilot testing and set to avoid ceiling and floor effects while achieving ~60-80% behavioral performance in both younger and older adults. Target speech was always presented at 75 dB_A_ SPL. Stimulus presentation was controlled through MATLAB (The MathWorks, Inc.; Natick, MA) via a TDT RP2 interface (Tucker-Davis Technologies; Alachua, FL). Stimuli were delivered binaurally via insert earphones (Etymotic Research; Elk Grove Village, IL).

**Figure 2.**
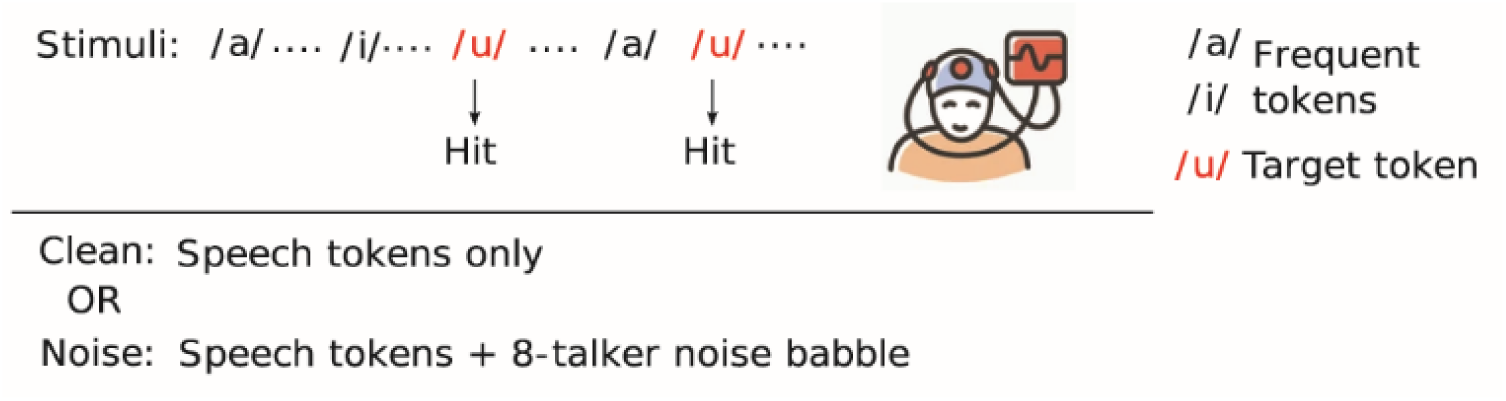
Example stimulus paradigm. Participants detected infrequent, oddball target tokens (ex. /u/) in a continuous stream of speech sounds presented at ~4.5 Hz. Participants identified targets via button press, and reaction time and hits/misses were logged. Stimuli were presented in clean and noise (+ 8-talker babble) conditions during simultaneous EEG acquisition. Figure adapted from Lai et al. (2022).

Participants were asked to respond via button press each time they detected the oddball target. Reaction time (RT) and signal detection accuracy (hits/misses) were logged for each trial. “Hits” were defined as the presence of a response within 5 token presentations of the oddball target (Price & Bidelman, 2021).

### 2.3 EEG recording and preprocessing

EEG pre-processing was conducted in MATLAB (The MathWorks, Inc., Natick, MA). EEGs were originally recorded from electrodes placed at 10-10 locations on the scalp (Oostenveld & Praamstra, 2001) and digitized > 5 kHz. However, the data were re-montaged to a common 37 channel electrode configuration (Fpz, Fp1/2, Fz, F3/4, F7/8, F9/10, FC5/6, FC1/2, T7/8, T9/10, Cz, C3/4, CP1/2, CP5/6, Pz, P3/4, P7/8, P9/10, Oz, O1/2, A1/A2) via interpolation (Desjardins, 2010) and downsampled to 250 Hz to equate channel counts and sampling rates across datasets. EEGs were re-referenced to linked mastoids (M1/M2) offline for subsequent analysis.

Continuous EEGs were then bandpass filtered from 2-50 Hz to isolate phase-locking to the speech envelope and epoched into 10-sec segments relative to target onsets (e.g., He et al., 2023). 500 epochs from each recording were then averaged to obtain evoked waveforms for each channel and equate the overall duration of data for PLV analyses across participants.

### 2.4 Electrophysiological data analysis

To assess brain-to-speech synchronization, we analyzed neural phase-locking value (PLV) to the speech stimulus (He et al., 2023; Lachaux et al., 1999). Procedures were similar to those described in He et al. (2023). PLV describes the correspondence between two signals by measuring the average phase consistency over time. To compute frequency-specific PLV, we first bandpass filtered the stimulus and EEG signals at each electrode around each frequency bin (±1 Hz) from 1 to 30 Hz (0.5 Hz steps). Signals were then windowed with 500 ms ramps to reduce onset/offset transients and the instantaneous phase was extracted using the Hilbert transform. PLV was then calculated in each narrow frequency band according to Equation 1:

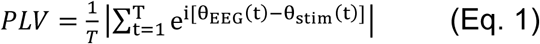

where *θ_EEG_* (*t*) and *θ_stim_* (*t*) are the instantaneous phases of the EEG and stimulus signals, respectively. PLV ranges from 0 to 1, where 0 represents no phase synchrony between signals, and 1 represents perfect phase synchrony. Repeating this procedure across frequencies (1–30 Hz; 0.5 Hz steps) resulted in a continuous function of PLV describing the degree of brain-to-speech synchronization across the bandwidth of interest (Assaneo et al., 2019; He et al., 2023). We then measured the peak PLV value between 4 and 6 Hz (i.e., the nominal rate of our speech stimuli) at each electrode channel to assess changes in speech phase-locking with age and SNR.

### 2.5 Behavioral data

To analyze behavioral responses in the speech detection task, we calculated signal detection metrics including *d*-prime (*d′*), hits (*H*), false alarm (*FA*) rate, and response bias (*c*) (Green & Swets, 1966; Macmillan & Creelman, 2005) to isolate listeners’ perceptual sensitivity from their decision criteria. A higher *d*′ [*d*′ = z(*H*) – z(*FA*)] represents greater sensitivity to detect oddballs within the continuous speech stream. Higher FA indicates more frequent errors when oddball tokens were not present but incorrectly reported as such. Higher bias (*c*) values (*c =* −0.5*[z(*H*) + z(*FA*)]) indicate a more conservative response (e.g., reduced tendency to respond “signal present” even when the oddball token was present). Cases where listeners obtain perfect accuracy (i.e., *H* = 1, *FA* = 0) imply a *d*′ of infinity. In these instances, hits and false alarms were corrected to 0.99 and 0.01, respectively, in order to compute finite signal detection metrics (Macmillan & Creelman, 2005).

### 2.6 Speech-in-noise perception

To assess relationships between our primary task and clinically-normed speech-in-noise perception, we administered two lists of the Quick Speech-in-Noise (QuickSIN) test (Killion et al., 2004). Participants repeat low-context sentences presented in increasing levels of background noise. This test results in a score reflecting SNR loss, reflecting the dB SNR for 50% keyword recall performance (0-3 dB SNR loss is considered normal).

### 2.7 Statistical analysis

Mixed-model ANOVAs were performed on the dependent variables (PLV, *d*′, FA, and c) in R (version 4.2.2, R-Core-Team 2020) using the lme4 package (Bates et al., 2015). Models utilized fixed predictors of group (2 levels; OA vs. YA) and SNR (2 levels; clean vs. noise), with subject-level random intercepts. Effect sizes are reported as partial eta squared 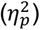 and degrees of freedom (*d.f.*) using Satterthwaite’s method. Tukey adjustments were used to correct for multiple pairwise comparisons.

We assessed relationships between behavioral, neural, and demographic variables using Spearman correlations (*r_s_*). For these analyses, we considered noise-related decrements in each neural and behavioral index (e.g., ΔPLV = PLV_clean_ – PLV_noise_; Δ*d*′ = *d*′_clean_ – *d*′_noise_). Change scores indexed the amount of degradation to brain and behavioral speech processing measures introduced by noise. PTA was positively correlated with age (*r* = 0.76, *p* < 0.0001; data not shown), indicating older adults had (expectedly) more hearing loss. Consequently, we partialled out PTA in the correlations to assess the impact of age on neural and behavioral measures independent of age-related presbycusis.

For the correlations between age vs. (i) neural (ΔPLV) and (ii) behavioral (Δ*d*′) variables, we also performed commonality analysis (Nimon & Reio, 2011). This decomposed the explained variance (*R^2^*) into variance shared between age and hearing thresholds (average bilateral thresholds, 250-8000 Hz), and unique variance contributed by these factors. We fit linear regression models including age and hearing threshold alone and together, separately. Unique variance for age and hearing threshold was determined by calculating the reduction in the overall model *R^2^*after removing each predictor from the model, and shared variance was calculated as the overlap between the predictions of age and hearing threshold.

## 3. Results

### 3.1 Behavioral results

Figure 3 displays behavioral results across groups and stimulus SNR. An ANOVA on *d*′ (Fig. 3A) revealed an interaction between group and SNR (*F*(1,81) = 11.72, *p* = 0.00097, 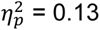). YA had greater decline in their perceptual sensitivity between clean and noise conditions (*t*(81) = 9.039, *p* < 0.0001) than did OA (*t*(81) = 2.93, *p* = 0.0044). A main effect of SNR was also present, with higher overall *d*′ scores in clean relative to noise conditions for both groups (*F*(1,81) = 63.61, *p* < 0.0001, 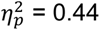).

**Figure 3.**
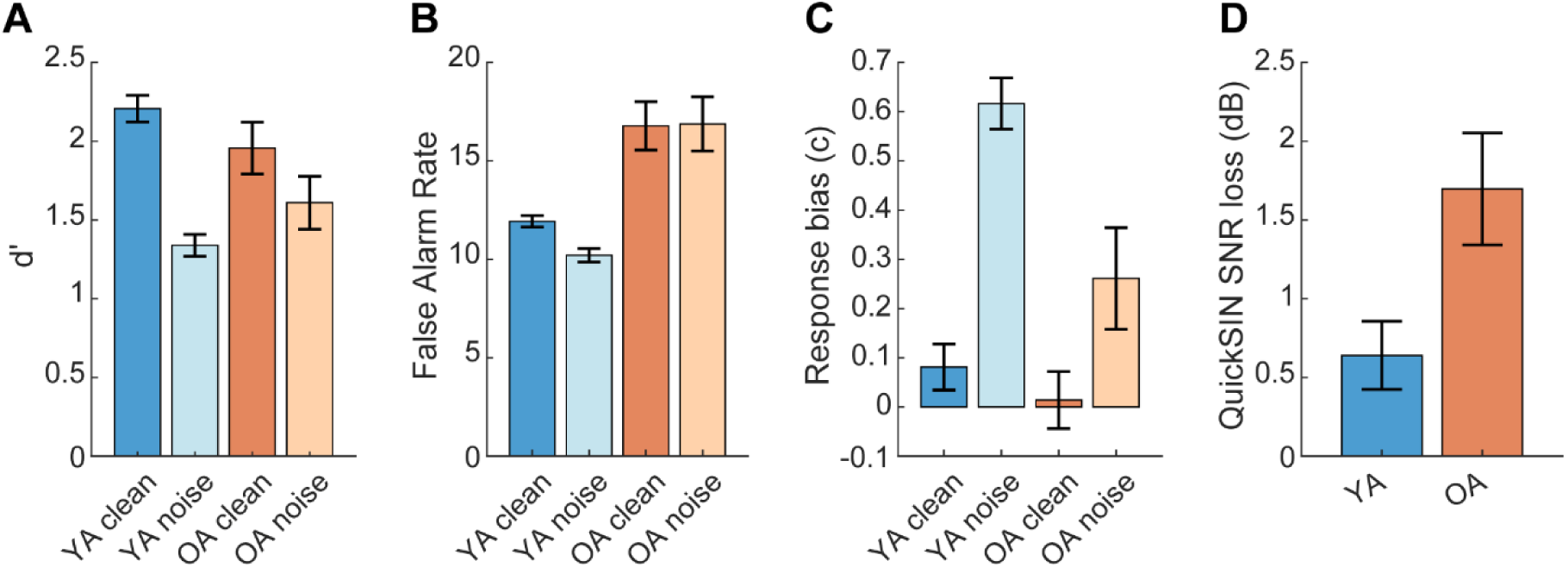
Behavioral results for syllable detection task. **(A)** YA had greater decline in *d*′ between clean and noise conditions than OA. **(B)** False alarm rates were higher for OA vs. YA. **(C)** YA had higher response bias (c) than OA in noise; groups showed similar bias for clean speech. **(D)** YA had lower (better) QuickSIN scores than OA. Error bars: ±1 S.E.M.

An ANOVA on FA rates (Fig. 3B) revealed a main effect of group. OA made more frequent false alarms than YA (*F*(1,81) = 35.84, *p* < 0.0001, 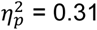). No other main effects or interactions were present (*p*s > 0.1).

An ANOVA on response bias (Fig. 3C) revealed an interaction between group and SNR (*F*(1,81) = 9.39, *p* = 0.0030, 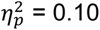), driven by more conservative responses for YA than OA, but only in noise (clean pairwise comparison: *t*(133) = −0.737, *p* = 0.46; noise: *t*(133) = −3.89, *p* = 0.0002). Main effects of group and condition confirmed more conservative response bias both for YA (*F*(1,81) = 7.32, *p* = 0.0083, 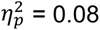) and for speech in noise (*F*(1,81) = 69.06, *p* < 0.0001, 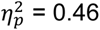), respectively.

Finally, a t-test on QuickSIN scores (Fig. 3D) revealed lower (better) scores in the YA group relative to OA (*t*(55.23) = 2.54, *p* = 0.014). This indicates older adults’ changes in SIN processing at the token level extend to poorer recognition performance at the sentence-level.

### 3.2 Neural speech tracking results

Figure 4 displays scalp topographies of peak PLV between 4-6 Hz, reflecting the degree of neural phase-locking to the ongoing speech stream. Difference maps revealed that OA had larger ΔPLV between clean and noise-degraded speech than YA. The distribution of PLV was generally strongest near the vertex (Cz), indicative of neural generators in the supratemporal plane, e.g., auditory cortex (Picton et al., 1999). Consequently, we further quantified PLV at the Cz electrode where these effects were most prominent over the scalp.

**Figure 4:**
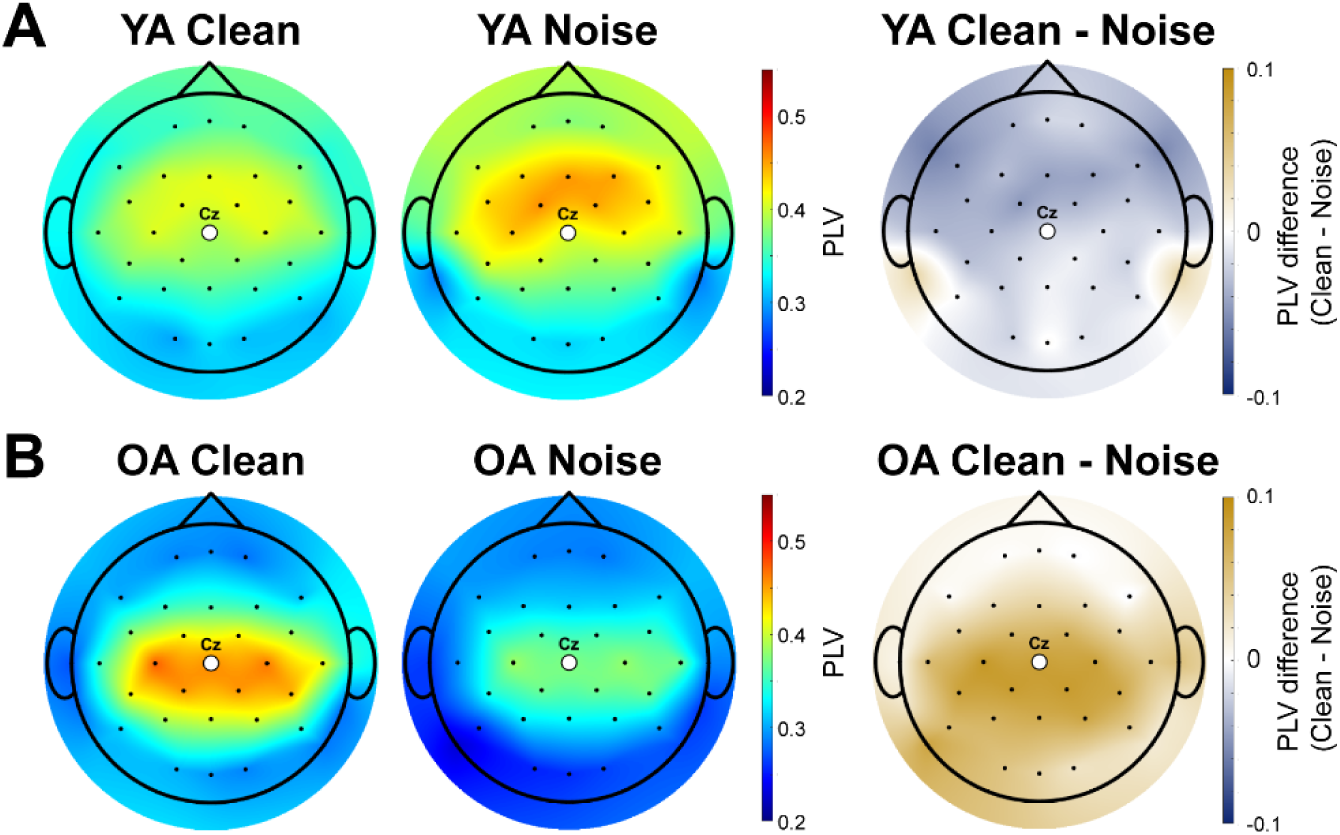
PLV topographies. Scalp maps show maximum PLVs between 4-6 Hz across the scalp for clean and noise-degraded speech for both **(A)** Younger and (**B**) Older Adults. Mean differences contrasting SNR conditions (Clean – Noise) per group are shown on right. Responses at Cz electrode (center, white circle) were used for statistical analyses.

Figure 5 shows PLV as a function of frequency per group and noise SNR. Note the peaks at ~5, 10, and 15 Hz, reflecting phase-locked energy to the fundamental and harmonic frequencies of the stimulus rate that is common in neural speech synchronization (cf. He et al., 2023).

**Figure 5.**
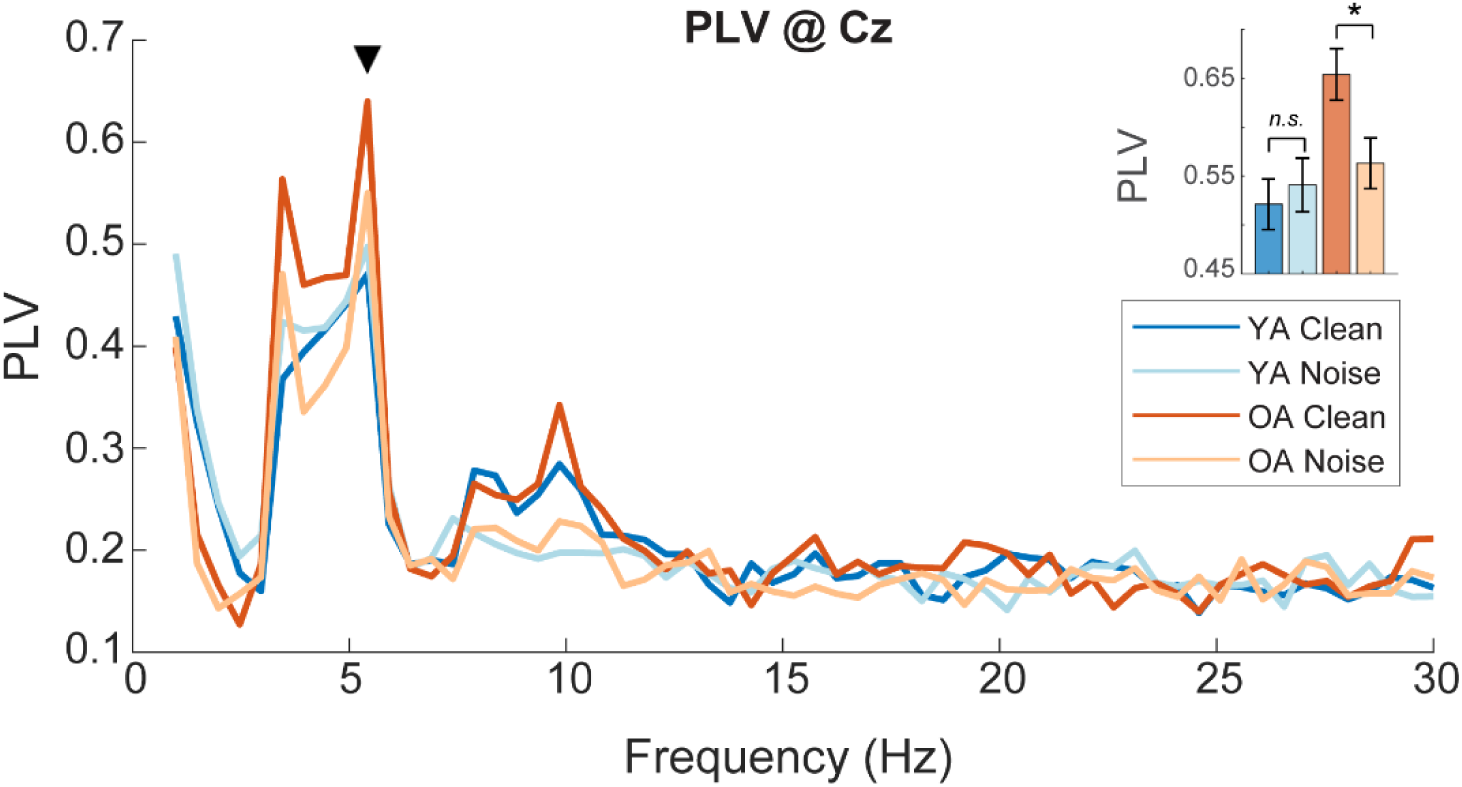
PLV reveals age-related changes in the neural speech tracking. PLV at the Cz electrode for each group and SNR condition across frequencies. Peak PLV at 4-6 Hz (▾), the nominal rate of the speech stimulus train. (*inset bars*) OA demonstrated significantly higher PLV to speech presented in clean vs. noise; YA had similar PLV regardless of speech SNR. Error bars: ±1 S.E.M.; \**p* < 0.05

An ANOVA on peak PLV between 4-6 Hz (▼, Fig. 5) revealed no main effect of SNR by itself (*p* = 0.13). However, there was a main effect of group (*F*(1,81) = 6.26, *p* = 0.014, 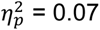), indicating higher overall PLV in older relative to younger listeners. More critically, we found a group x SNR interaction (*F*(1,81) = 5.40, *p* = 0.023, 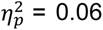). This interaction was driven by higher PLVs in OA for clean relative to noisy speech (pairwise contrast: *t*(81) = 2.46, *p* = 0.016). In contrast, YA had similar PLV to both clean and noisy speech (*t*(81) = −0.66, *p* = 0.51).

### 3.4 Relationships between oddball token detection, neural synchronization, and demographic variables

Figure 6 shows correlations between demographic (age, hearing thresholds), neural (ΔPLV), and behavioral (Δd′) measures. We partialled out hearing thresholds (averaged bilateral thresholds 250-8000 Hz) in these correlations due to the strong association between hearing loss and age in our sample (see *Sect. 2.6*). After controlling for hearing, age became a much weaker predictor of ΔPLV (Fig. 6A; before partial: *r* = 0.34, *p* = 0.0015; after partial: *r* = 0.183, *p* = 0.099) but remained a strong predictor of Δ*d′* (Fig. 6B; before partial: *r* = −0.36, *p* = 0.00077; after partial: *r* = −0.247, *p* = 0.025). That is, older listeners experienced smaller perceptual decrements in speech processing with increasing noise. There were no relationships between brain (neural ΔPLV) and behavior (Δ*d′*) in the EEG task (*r* = 0.032, *p =* 0.77, Fig. 6D). However, hearing thresholds did correlate with QuickSIN scores and remained significant even when controlling for age (before partial: *r* = 0.33, *p* 0.0025; after partial: *r* = 0.33, *p* = 0.0020, Fig. 6E).

**Figure 6.**
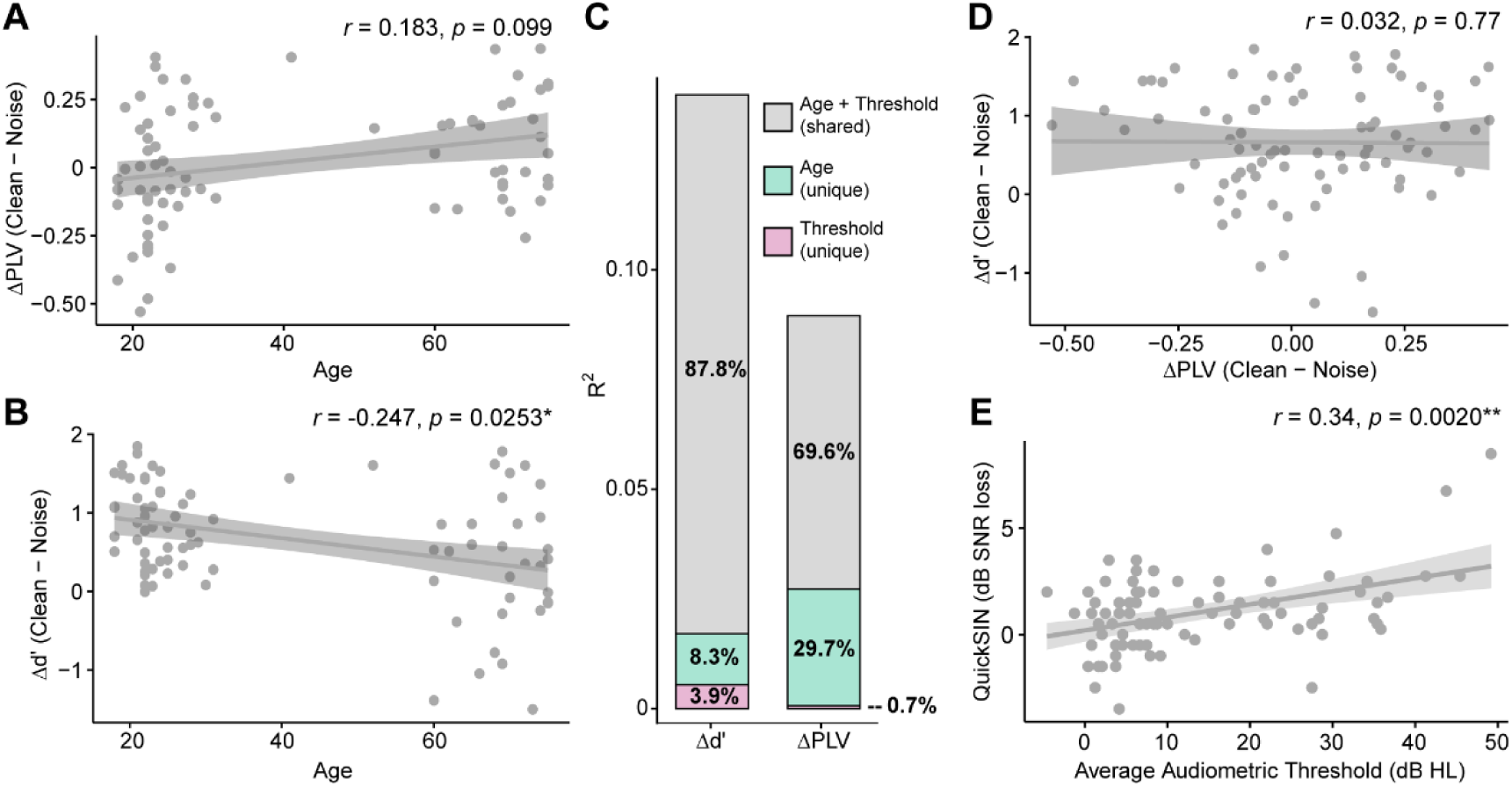
Age influences neurobehavioral speech tracking even when controlling for hearing status. Scatters show age vs. (**A**) neural ΔPLV and (**B**) perceptual Δ*d′* between clean and noise-degraded speech. After controlling for average hearing thresholds across the audiogram, the relationship between age and noise-related degradations to PLV is no longer significant (*p* = 0.099), but the relationship between age and noise-related degradation in *d′* remains (*p* < 0.05). **(C)** Commonality analysis reveals the unique contribution of age and hearing to variance explained. Shared variance between age and hearing accounts for the majority of variance (*R^2^*) in neuro-behavioral measures. (**D**) No relation between ΔPLV and Δ*d′*. (**E**) Audiometric thresholds positively correlate with QuickSIN SNR loss even after controlling for age. \**p* < 0.05, \*\**p* < 0.01, shading = 95% CI.

To understand the relative contributions of age and hearing to changes in neural and behavioral outcomes, we performed a commonality analysis (Nimon & Reio, 2011) using standard linear multiple regression models (Fig. 6C). For behavioral Δ*d′* (total *R^2^* = 0.140), the shared contribution between age and hearing thresholds explained a large majority of variance explained by our model (*R^2^_age+hearing_* = 0.123, 87.8%), followed by smaller contributions from age alone (*R^2^_age_* = 0.012, 8.3%) and hearing alone (*R^2^_hearing_* = 0.123, 87.8%). For neural ΔPLV (total *R^2^* = 0.0900), shared contributions from age and hearing also contributed to a large proportion of the total explained variance (*R^2^_age+hearing_* = 0.062, 69.6%), followed by age alone (*R^2^_age_* = 0.027, 29.7%), and finally a very small contribution from hearing alone (*R^2^_hearing_* = 0.001, 0.7%).

### 3.5 Model EEG simulations of PLV

To assess whether PLV effects in our empirical data might be explained by more trivial differences in stimulus acoustics, we simulated oscillatory EEG responses to our speech train stimuli using a simple convolution model of evoked potential generation (e.g., Goldstein & Kiang, 1958; Janssen et al., 1991). The model is based on the assumption that sustained auditory evoked potentials are a series of overlapping onset evoked responses generated by repeating events in a running stimulus (e.g., Bidelman, 2015; Bidelman et al., 2024; Daly et al., 1976; Dau, 2003; Davis & Hirsh, 1974; Galambos et al., 1981; Gerken et al., 1975; Janssen et al., 1991; Picton et al., 1977). The approach is similar to that described in our previous papers (Bidelman, 2015; Bidelman et al., 2024) and is akin to the temporal response function (TRF) analysis (Crosse et al., 2021) recently popularized in the field.

EEGs were simulated by convolving a unitary response (“kernel”) waveform with the continuous acoustic envelope of the speech signal (Fig. 7A). We modelled the unitary TRF describing the canonical auditory cortical potentials (i.e., “P1-N1-P2” complex) as a sum of Gaussian functions (Equation 2):

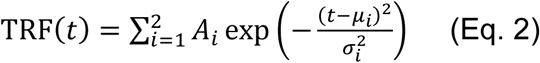

where 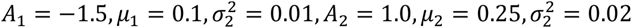. Convolving the transient wavelet [*TRF(t)*] with the acoustic envelope [*Stim(t)*] resulted in a complex waveform simulating the EEG to the continuous speech train [i.e., *EEG_model_(t) = TRF(t) * Stim(t)*]. Both white (Gaussian) and alpha-band noise (8-12 Hz, 2^nd^ order Butterworth) were then added to model signals at √10 and 1.5 amplitude, respectively, to simulate realistic EEG noise.

**Figure 7:**
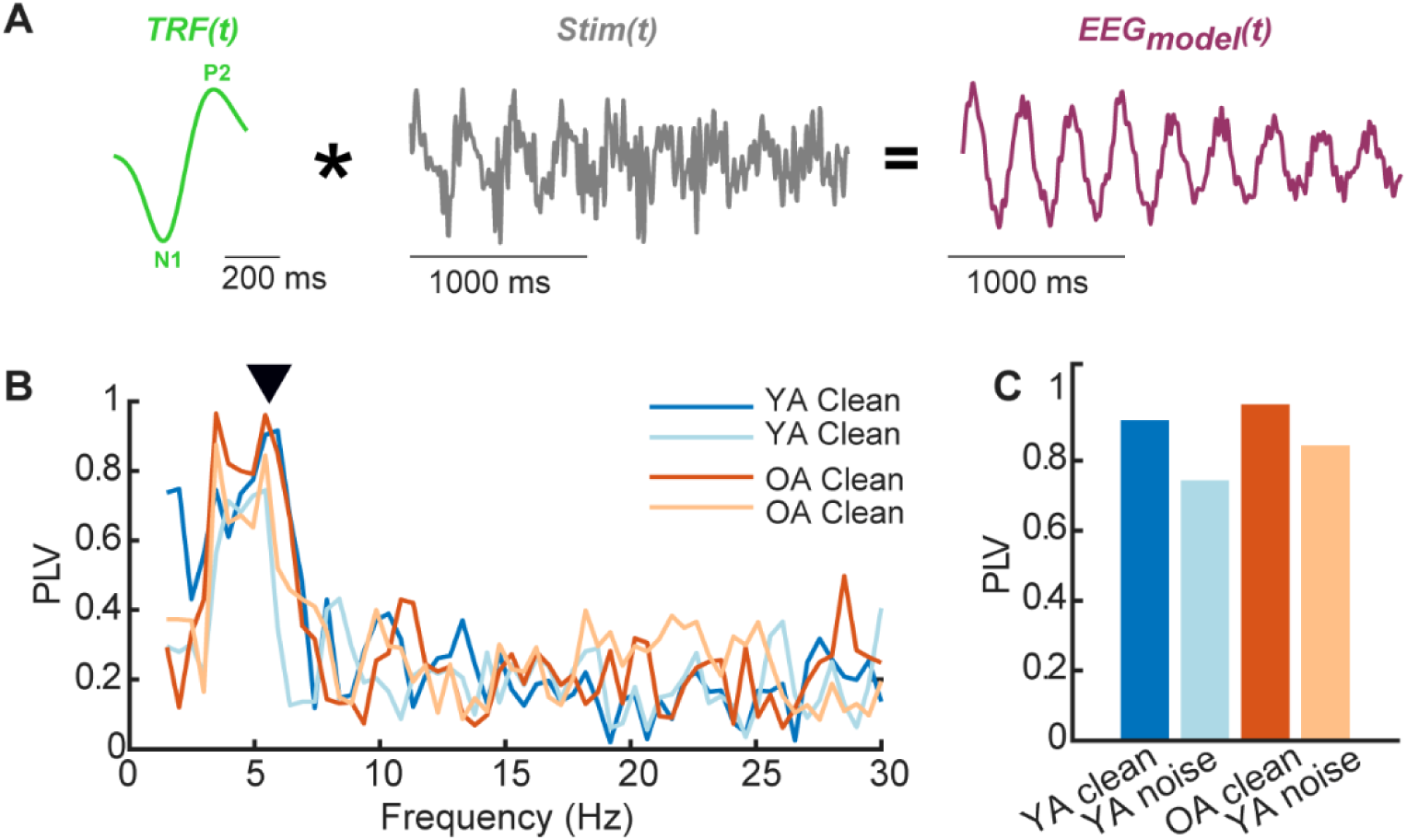
Simulated PLV from model EEG responses. **(A)** EEGs to speech trains are modeled as a convolution between the unit TRF response (cf. canonical P1-N1-P2) and the acoustic speech envelope of the continuous speech train (for details, see Bidelman, 2015; Bidelman et al., 2024). The resulting complex waveform simulates a sustained evoked potential to a periodic stimulus. (**B**) PLV computed from model responses using vowel vs. CV trains and +5 and +10 dB SNR simulating EEG data for “younger” and “older” adults according to original dataset parameters. Compare to actual EEG recordings in Fig. 5. (**C**) Peak PLV simulations for clean and noise-degraded speech. Model data diverge from empirical results suggesting group differences in PLV (cf. Fig. 5) cannot be explained by differences in stimulus acoustics.

Model responses were then analyzed using identical procedures as in the empirical recordings including filter, epoch, and stimulus used for PLV calculations (Fig. 7B, C; see *Sect. 2.4*). That is, PLV was computed between the simulated EEG responses and the evoking stimulus using the respective tokens (vowels, CVs) and SNRs (+5 dB, +10 dB) as used in the corresponding original studies (Bidelman, Mahmud, et al., 2019; Bidelman, Price, et al., 2019; Bidelman et al., 2026; Price & Bidelman, 2021). Divergence in the pattern of model and actual data would suggest group differences in PLV are not due solely to exact token or SNR choice but instead reflect additional age- and/or hearing-related changes in neural phase locking beyond those explainable by mere acoustic differences.

## 4. Discussion

Using EEG, we assessed how aging impacts neural synchronization to and identification of target speech in clean and noise-degraded listening conditions. We found that: (a) older adults (OA) demonstrated fewer behavioral differences between clean and noise than did younger adults (YA); (b) OA demonstrated greater noise-related reductions in neural speech tracking speech than YA, despite having overexaggerated neural tracking of clean speech; and (c) age accounted for behavioral (but not neural) deficits in speech processing beyond the effects of hearing loss. Lastly, computational modeling suggested these age-related differences are not due to acoustic stimulus properties alone but instead reflect changes in the neurobiological synchronization to complex sounds with advancing age.

### 4.1 Aging is associated with less behavioral sensitivity to noise-related changes

In a speech-oddball detection paradigm, YA showed expected noise decrements in perception whereas OA performed similarly across SNRs (i.e., Δ*d′*_OA_ < Δ*d′*_YA_). The more subtle behavioral changes seen in OA aligns with the reduced release from masking and gap listening observed in elderly listeners (Dubno et al., 2002; Feng et al., 2018; Irsik et al., 2022; Zobel et al., 2019). OA also had higher overall false alarm (FA) rates across listening conditions and were less conservative responders (lower response bias) than YA. Older adults might rely on a more widely dispersed or inefficient neural network during speech perception (Voytek & Knight, 2015; Wong et al., 2009), which may contribute to increased internal noise and therefore a greater tendency to falsely respond even when no signal was present (Salthouse & Lichty, 1985) (but see (Bidelman et al., 2014; Voytek et al., 2015). These mechanisms could contribute to the changes we find in older listeners’ perceptual performance.

Speech perception deficits in OA are thought to arise from a complex, dynamic interplay between lower-level sensory/perceptual decline and higher-level cognitive processing (Schneider et al., 2002). Aging is also associated with greater reliance on linguistic-semantic contextual information to compensate for decreased sensory input (Gordon-Salant & Fitzgibbons, 1997; Nittrouer & Boothroyd, 1990). Thus, the lesser noise-related change we find in older adults’ behavior may reflect an increased reliance on top-down processing to offset a more impoverished and error-prone acoustic signal. While increased top-down processing can offset age-related declines in SIN perception (Irsik et al., 2022), it comes at the cost of cognitive efficiency (Wingfield & Tun, 2001; Wingfield et al., 2005), and at times can lead to “false hearing” (Rogers et al., 2012). More “false hearing” could account for the increased bias and false alarms we find in older listeners’ speech detection. However, our continuous speech stream paradigm did not contain contextual cues so top-down compensation cannot be due to linguistic processing, per se. Alternatively, older adults may exert more listening effortful in noise (Doherty & Desjardins, 2013; Gosselin & Gagne, 2011), which could explain the lesser age-related change in Δ*d′* we find in those participants.

### 4.2 Age-related differences in auditory neural synchronization to speech

#### 4.2.1. Older adults show overexaggerated PLV to clean speech

Contrasting the behavioral results, OA demonstrated overexaggerated neural synchronization (PLV) to syllable trains presented in clean, and consequently, greater reductions in PLV with noise. Electrophysiological and psychoacoustic findings in both animals and humans show that aging and hearing loss are associated with increased envelope coding (Bacon & Gleitman, 1992; Kale & Heinz, 2010; Moore & Glasberg, 2001; Parthasarathy et al., 2014; Schmitt et al., 2022). Aging and ARHL are inextricably tied and often manifest in overlapping effects (see *Sect. 4.3.*). One possibility is that stronger neural responses in elderly individuals could reflect “central gain” mechanisms (Chambers et al., 2016) which help offset the reduced sensory input (Millman et al., 2017) associated with auditory aging. Such overexaggerated envelope encoding could reflect a general imbalance between excitatory/inhibitory processes in the central auditory nervous system (e.g., Caspary et al., 2008; Caspary et al., 1995). Alternatively, increased neural tracking of speech in older adults may be related to increased listening effort (Decruy et al., 2019), though the influence of attentional effort on neural tracking changes with speech intelligibility (He et al., 2025). Our scalp EEG data cannot pinpoint the exact mechanism behind age-related envelope enhancement. However, our findings do support work suggesting that greater neural envelope coding of syllabic-rate modulations (i.e., 2-5 Hz) may not always be perceptually advantageous for robust speech intelligibility (Goossens et al., 2018; Millman et al., 2017). It is also possible that naturalistic speech processing relies on several different neural mechanisms including (but certainly not limited to) auditory neural synchronization.

With regard to hearing acuity, enhanced envelope responses in older adults could result from a loss of cochlear compression, loudness recruitment, and broadening of the auditory filters with peripheral hearing loss (Wallaert et al., 2017). ARHL is also associated with hyperactive midbrain and cortical evoked responses to speech (Bidelman et al., 2014) and enhanced envelope representation (Goossens et al., 2018; Herrmann, 2025; Presacco et al., 2016b). Additionally, in hearing-impaired listeners, larger speech comprehension benefits scale with neural speech tracking accuracy as measured via EEG (Schmitt et al., 2022) (but see Petersen et al., 2017). Our PLV data are largely compatible with the decline-compensation hypothesis of aging (Wong et al., 2009), showing unusually large acoustic envelope representation in brain responses of aging adults.

#### 4.2.2. Older adults demonstrated greater noise-related degradation in neural speech tracking

Despite stronger neural tracking of clean speech, OA experienced greater reductions in neural tracking with noise compared to YA. Previous studies have also observed exaggerated speech envelope encoding paired with greater noise-related reductions in older adults (Presacco et al., 2016b). Such effects are especially evident when the background noise is quasi-intelligible (Presacco et al., 2016a), as in our paradigm with speech babble. Degradations to SIN processing with age are also attributed to deficits in fine structure (Anderson et al., 2013; Grose & Mamo, 2010; Strelcyk & Dau, 2009) and temporal processing (Anderson et al., 2020; Gordon-Salant et al., 2011), perceptual attributes which are easily degraded by background noise.

Hearing loss and aging both impact temporal processing (McClaskey, 2024) and are highly intercorrelated factors in auditory aging. Yet, associations between broader temporal processing deficits and speech perception persist even when adjusting for hearing and cognitive ability (Babkoff & Fostick, 2017). Though our noise stimuli were not titrated based on individual performance, and the SNR differed slightly across datasets, it is notable we observe differences in OA neural synchronization even to clean speech envelopes. Similar results have been observed in recent studies in both geriatric (Herrmann, 2025) and pediatric listeners who present with central auditory processing deficits (Momtaz et al., 2021). Moreover, our modeling results predicted equally strong neural phase-locking to clean speech in younger and older ears and smaller noise-related changes in elderly listeners. Yet, this is not what we observe in our data. This suggests the observed differences in speech phase-locking between younger and older listeners are not due to subtle or differences in speech acoustics alone, but rather, reflect more salient neurobiological changes in complex auditory processing due to the various factors of aging.

### 4.3 Relationships between age, hearing loss, and speech tracking

As expected, we observed a strong relationship between age and hearing loss in nearly all of our speech measures. Though age and hearing are sometimes separable in describing speech processing in older adults (Babkoff & Fostick, 2017; Bidelman et al., 2014; Füllgrabe et al., 2015), our data confirm the multidimensional and highly interconnected nature of factors that impact auditory aging (Bugannim et al., 2025; Humes et al., 2013; Humes et al., 2012; Lentz et al., 2022). Here, we show that after partialling for hearing acuity, the relationship between age and neural ΔPLV disappears. Similarly, age considered jointly with hearing loss contributed substantial shared variance that explained EEG and perceptual responses and had more explanatory power than when considered separately. Thus, age by itself did not uniquely explain changes in neural tracking. In contrast, we found that the relationship between age and perceptual Δ*d′* remained significant even after controlling for hearing loss. These results together suggest that age was a unique (albeit small) predictor of noise-related differences in behavioral speech processing. In their systematic review specifically on speech tracking, Ratelle and Tremblay (2025) similarly conclude that both age and hearing acuity interdependently modulate neural synchronization. Though we did not screen for cognitive functioning, cognition is unlikely to account for our group effects as speech envelope tracking does not vary with cognitive status (Bolt & Giroud, 2024).

Our data highlight the importance of considering both age and ARHL when assessing speech synchronization in older listeners. This is particularly important for studies attempting to enhance speech envelope tracking and improve SIN processing via neurostimulation techniques (e.g., Erkens et al., 2020; Guilleminot & Reichenbach, 2022; van Bree et al., 2021; Wilsch et al., 2018). The present results suggest speech envelope tracking in older listeners is already hyperactive in a baseline state. When hearing acuity is considered, some studies find that age and ARHL uniquely contribute to differences in neural speech tracking (Bidelman et al., 2014; McClaskey, 2024; Petersen et al., 2017), while others do not (McClaskey et al., 2019). It is possible that discrepancies across studies could be in part due to stimulus audibility. When hearing thresholds are not well-controlled or accounted for, the relative contributions of aging and presbycusis can be confounded in speech tracking measures. The amount of shared variance between age and hearing loss seen in our neural PLV data underscores the importance of considering both hearing thresholds and age when studying senescent changes in neuroacoustic speech tracking.

Unfortunately, we did not detect a relationship between neural (PLV) and behavioral (*d′*) measures in our primary EEG task. This contrasts previous studies reporting an association between neural-speech PLV and SIN perception (Mai et al., 2018; McClaskey et al., 2019). Additionally, there was no relationship between QuickSIN scores and our EEG-based SIN task (but see Lai et al., 2022), despite the overall poorer QuickSIN scores in older vs. younger adults. The lack of correspondence between tasks suggests SIN perception may rely on a more complex set of processes than auditory synchronization alone. This discordance is especially true for the QuickSIN which requires sentence level recognition and likely taps high-order processes (e.g., working memory) that are probably not recruited during our simplistic EEG speech detection task.

### 4.4 Ramifications for temporal cuing interventions to improve SIN in older listeners

OA demonstrated less behavioral distinction between speech presented in clean vs. noise, greater noise-related degradations in neural synchronization to speech, and poorer SIN perception as assessed via the QuickSIN. Though enhanced envelope encoding is often assumed to improve speech perception (Decruy et al., 2019; Vanthornhout et al., 2018), the overexaggerated responses we find in older listeners (even for even quiet speech) tempers this assertion. Still, SIN deficits and poorer auditory temporal processing are among the most common hearing deficits in OA (Anderson et al., 2012; Füllgrabe et al., 2015; Gordon-Salant et al., 2011). Consequently, our data prompt questions on whether rehabilitation strategies to bolster temporal envelope processing might be beneficial in older individuals. Indeed, promising studies suggest that timing-based rate discrimination training can partially restore age-related deficits in temporal processing (Anderson et al., 2022). Cueing the speech envelope has also been shown to improve neural tracking to and perception of SIN in young adults (Falk et al., 2017; te Rietmolen et al., 2025). Relatedly, neurostimulation (e.g., tDACs, TMS) and multimodal stimuli (e.g., audio-haptic presentation) that aim to enhance envelope tracking by mirroring sound with other concurrent signals are also encouraging (e.g., Erkens et al., 2020; Guilleminot & Reichenbach, 2022; van Bree et al., 2021; Wilsch et al., 2018). However, putative benefits have not yet been widely explored in older listeners. It is possible that temporal cueing of SIN could minimize the greater amount of noise-related degradation seen in speech synchronization in older ears. Alternatively, there may be a fundamental limit on to the degree to which speech envelope enhancement remains viable for perception in older listeners, as suggested by our data. Future studies should investigate the potential benefits and limits of temporal cueing to restore envelope encoding in the aging auditory system.

## Funding

This work was supported by a grant from the National Institutes of Health (NIA) awarded to J.M. (1F31AG101836).

## Notes

### Competing Interest Statement

The authors have declared no competing interest.

## References

1. Anderson, S., DeVries, L., Smith, E., Goupell, M. J., & Gordon-Salant, S. (2022). Rate Discrimination Training May Partially Restore Temporal Processing Abilities from Age-Related Deficits. Journal of the Association for Research in Otolaryngology, 23(6), 771–786. 10.1007/s10162-022-00859-x

2. Anderson, S., Parbery-Clark, A., White-Schwoch, T., Drehobl, S., & Kraus, N. (2013). Effects of hearing loss on the subcortical representation of speech cues. Journal of the Acoustical Society of America, 133(5), 3030–3038. 10.1121/1.4799804

3. Anderson, S., Parbery-Clark, A., White-Schwoch, T., & Kraus, N. (2012). Aging affects neural precision of speech encoding. Journal of Neuroscience, 32(41), 14156–14164. 10.1523/jneurosci.2176-12.2012

4. Anderson, S., Roque, L., Gaskins, C. R., Gordon-Salant, S., & Goupell, M. J. (2020). Age-related compensation mechanism revealed in the cortical representation of degraded speech. Journal of the Association for Research in Otolaryngology, 21(4), 373–391. 10.1007/s10162-020-00753-4

5. Assaneo, M. F., & Poeppel, D. (2018). The coupling between auditory and motor cortices is rate-restricted: Evidence for an intrinsic speech-motor rhythm. Sciences Advances, 4(2), eaao3842. 10.1126/sciadv.aao3842

6. Assaneo, M. F., Ripollés, P., Orpella, J., Lin, W. M., de Diego-Balaguer, R., & Poeppel, D. (2019). Spontaneous synchronization to speech reveals neural mechanisms facilitating language learning. Nature Neuroscience, 22, 627–632. 10.1038/s41593-019-0353-z

7. Babkoff, H., & Fostick, L. (2017). Age-related changes in auditory processing and speech perception: cross-sectional and longitudinal analyses. European Journal of Ageing, 14(3), 269–281. 10.1007/s10433-017-0410-y

8. Bacon, S. P., & Gleitman, R. M. (1992). Modulation detection in subjects with relatively flat hearing losses. Journal of Speech, Language, and Hearing Research, 35(3), 642–653. 10.1044/jshr.3503.642

9. Bates, D., Mächler, M., Bolker, B., & Walker, S. (2015). Fitting linear mixed-effects models using lme4. Journal of Statistical Software, 67(1), 1–48. 10.18637/jss.v067.i01

10. Beck, D., Larsen, D., & Bush, E. (2018). Speech in noise: Hearing loss, neurocognitive disorders, aging, traumatic brain injury and more. Journal of Otolaryngology ENT Research, 10(4), 199–205. 10.15406/joentr.2018.10.00345

11. Bidelman, G. M. (2015). Multichannel recordings of the human brainstem frequency-following response: Scalp topography, source generators, and distinctions from the transient ABR. Hearing Research, 323, 68–80. 10.1016/j.heares.2015.01.011

12. Bidelman, G. M., Lowther, J. E., Tak, S. H., & Alain, C. (2017). Mild cognitive impairment is characterized by deficient hierarchical speech coding between auditory brainstem and cortex. Journal of Neuroscience, 37(13), 3610–3620.

13. Bidelman, G. M., Mahmud, M. S., Yeasin, M., Shen, D., Arnott, S., & Alain, C. (2019). Age-related hearing loss increases full-brain connectivity while reversing directed signaling within the dorsal-ventral pathway for speech. Brain Structure and Function, 224(8), 2661–2676. 10.1007/s00429-019-01922-9

14. Bidelman, G. M., Price, C. N., Shen, D., Arnott, S. R., & Alain, C. (2019). Afferent-efferent connectivity between auditory brainstem and cortex accounts for poorer speech-in-noise comprehension in older adults. Hearing Research, 382(107795), 1–12. 10.1016/j.heares.2019.107795

15. Bidelman, G. M., Sisson, A., Rizzi, R., MacLean, J., & Baer, K. (2024). Myogenic artifacts masquerade as neuroplasticity in the auditory frequency-following response. Frontiers in Neuroscience, 18 - 2024. 10.3389/fnins.2024.1422903

16. Bidelman, G. M., Stirn, J. R., Rizzi, R., MacLean, J. A., & Cheng, H. (2026). Auditory brainstem–cortical anatomy relates to the magnitude of frequency-following responses (FFRs) and event-related potentials (ERPs) coding speech-in-noise. Neuroimaging, 1(1), 6. 10.3390/neuroimaging1010006

17. Bidelman, G. M., Villafuerte, J. W., Moreno, S., & Alain, C. (2014). Age-related changes in the subcortical-cortical encoding and categorical perception of speech. Neurobiology of Aging, 35(11), 2526–2540. 10.1016/j.neurobiolaging.2014.05.006

18. Bolt, E., & Giroud, N. (2024). Auditory encoding of natural speech at subcortical and cortical levels is not indicative of cognitive decline. eNeuro, 11(5), 1–15. 10.1523/eneuro.0545-23.2024

19. Bugannim, Y., Roziner, I., & Kishon-Rabin, L. (2025). Speech recognition in noise across the life span with cognition and hearing sensitivity as mediators of age effects. Scientific Reports, 15(1), 20575. 10.1038/s41598-025-05882-5

20. Caspary, D. M., Ling, L., Turner, J. G., & Hughes, L. F. (2008). Inhibitory neurotransmission, plasticity and aging in the mammalian central auditory system. Journal of Experimental Biology and Medicine, 211, 1781–1791. 10.1242/jeb.013581

21. Caspary, D. M., Milbrandt, J. C., & Helfert, R. H. (1995). Central auditory aging: GABA changes in the inferior colliculus. Experimental Gerontology, 30(3), 349–360. 10.1016/0531-5565(94)00052-5

22. Chambers, A. R., Resnik, J., Yuan, Y., Whitton, J. P., Edge, A. S., Liberman, M. C., & Polley, D. B. (2016). Central gain restores auditory processing following near-complete cochlear denervation. Neuron, 89(4), 867–879. 10.1016/j.neuron.2015.12.041

23. Crosse, M. J., Zuk, N. J., Di Liberto, G. M., Nidiffer, A. R., Molholm, S., & Lalor, E. C. (2021). Linear modeling of neurophysiological responses to speech and other continuous stimuli: Methodological considerations for applied research. Frontiers in Neuroscience, 15(1350). 10.3389/fnins.2021.705621

24. Cruickshanks, K. J., Wiley, T. L., Tweed, T. S., Klein, B. E., Klein, R., Mares-Perlman, J. A., & Nondahl, D. M. (1998). Prevalence of hearing loss in older adults in Beaver Dam, Wisconsin. The epidemiology of hearing loss study. American Journal of Epidemiology, 148(9), 879–886. 10.1093/oxfordjournals.aje.a009713

25. Daly, D. M., Roeser, R. J., & Moushegian, G. (1976). The frequency-following response in subjects with profound unilateral hearing loss. Electroencephalography and Clinical Neurophysiology, 40, 132–142. 10.1016/0013-4694(76)90158-9

26. Dau, T. (2003). The importance of cochlear processing for the formation of auditory brainstem and frequency following responses. Journal of the Acoustical Society of America, 113(2), 936–950. 10.1121/1.1534833

27. Davis, H., & Hirsh, S. K. (1974). Interpretation of the human frequency following response. Journal of the Acoustical Society of America, 56, S63. 10.1121/1.1914281

28. Decruy, L., Vanthornhout, J., & Francart, T. (2019). Evidence for enhanced neural tracking of the speech envelope underlying age-related speech-in-noise difficulties. Journal of Neurophysiology, 122(2), 601–615. 10.1152/jn.00687.2018

29. Desjardins, J. A. (2010). Interpmont: EEGLAB Extension for Interpolating Recording Locations [Software]. GitHub. https://github.com/jadesjardins/interp_mont

30. Ding, N., Chatterjee, M., & Simon, J. Z. (2014). Robust cortical entrainment to the speech envelope relies on the spectro-temporal fine structure. Neuroimage, 88, 41–46. 10.1016/j.neuroimage.2013.10.054

31. Doherty, K., & Desjardins, J. A. (2013). Age-related changes in listening effort for various types of masker noises. Ear and Hearing, 34(3), 261–272. 10.1097/AUD.0b013e31826d0ba4

32. Dubno, J. R., Dirks, D. D., & Morgan, D. E. (1984). Effects of age and mild hearing loss on speech recognition in noise. The Journal of the Acoustical Society of America, 76(1), 87–96. 10.1121/1.391011

33. Dubno, J. R., Horwitz, A. R., & Ahlstrom, J. B. (2002). Benefit of modulated maskers for speech recognition by younger and older adults with normal hearing. The Journal of the Acoustical Society of America, 111(6), 2897–2907. 10.1121/1.1480421

34. Erkens, J., Schulte, M., Vormann, M., & Herrmann, C. S. (2020). Lacking effects of envelope transcranial alternating current stimulation indicate the need to revise envelope transcranial alternating current stimulation methods. Neuroscience Insights, 15, 2633105520936623. 10.1177/2633105520936623

35. Erkens, J., Schulte, M., Vormann, M., Wilsch, A., & Herrmann, C. S. (2021). Hearing impaired participants improve more under envelope-transcranial alternating current stimulation when signal to noise ratio is high. Neuroscience Insights, 16, 2633105520988854. 10.1177/2633105520988854

36. Etard, O., & Reichenbach, T. (2019). Neural speech tracking in the theta and in the delta frequency band differentially encode clarity and comprehension of speech in noise. The Journal of Neuroscience, 39(29), 5750–5759. 10.1523/jneurosci.1828-18.2019

37. Falk, S., Lanzilotti, C., & Schön, D. (2017). Tuning neural phase entrainment to speech. Journal of Cognitive Neuroscience, 29(8), 1378–1389. 10.1162/jocn_a_01136

38. Feng, T., Chen, Q., & Xiao, Z. (2018). Age-related differences in the effects of masker cuing on releasing Chinese speech from informational masking. Frontiers in Psychology, 9, 1–16. 10.3389/fpsyg.2018.01922

39. Füllgrabe, C., Moore, B. C. J., & Stone, M. A. (2015). Age-group differences in speech identification despite matched audiometrically normal hearing: contributions from auditory temporal processing and cognition. Frontiers in Aging Neuroscience, 6. 10.3389/fnagi.2014.00347

40. Galambos, R., Makeig, S., & Talmachoff, P. J. (1981). A 40-Hz auditory potential recorded from the human scalp. Proceedings of the National Academy of Sciences of the United States of America, 78(4), 2643–2647. 10.1073/pnas.78.4.2643

41. Gerken, G. M., Moushegian, G., Stillman, R. D., & Rupert, A. L. (1975). Human frequency-following responses to monoaural and binaural stimuli. Electroencephalography and Clinical Neurophysiology, 38, 379–386. 10.1016/0013-4694(75)90262-x.

42. Gifford, R. H., Bacon, S. P., & Williams, E. J. (2007). An examination of speech recognition in a modulated background and of forward masking in younger and older listeners. Journal of Speech, Language, and Hearing Research, 50(4), 857–864. 10.1044/1092-4388(2007/060)

43. Goldstein, M. H., & Kiang, N. Y. S. (1958). Synchrony of neural activity in electric responses evoked by transient acoustic stimuli. Journal of the Acoustical Society of America, 30, 107–114. 10.1121/1.1909497

44. Goossens, T., Vercammen, C., Wouters, J., & van Wieringen, A. (2018). Neural envelope encoding predicts speech perception performance for normal-hearing and hearing-impaired adults. Hearing Research, 370, 189–200. 10.1016/j.heares.2018.07.012

45. Gordon-Salant, S., & Fitzgibbons, P. J. (1997). Selected cognitive factors and speech recognition performance among young and elderly listeners. Journal of Speech, Language, and Hearing Research, 40(2), 423–431. 10.1044/jslhr.4002.423

46. Gordon-Salant, S., Fitzgibbons, P. J., & Yeni-Komshian, G. H. (2011). Auditory temporal processing and aging: Implications for speech understanding of older people. Audiology Research, 1(e4), 9–15. 10.4081/audiores.2011.e4

47. Gosselin, P. A., & Gagne, J. P. (2011). Older adults expend more listening effort than young adults recognizing speech in noise. Journal of Speech, Language, and Hearing Research, 54(3), 944–958. 10.1044/1092-4388(2010/10-0069)

48. Green, D. M., & Swets, J. A. (1966). Signal detection theory and psychophysics. Wiley.

49. Gregan, M. J., Nelson, P. B., & Oxenham, A. J. (2013). Behavioral measures of cochlear compression and temporal resolution as predictors of speech masking release in hearing-impaired listeners. The Journal of the Acoustical Society of America, 134(4), 2895–2912. 10.1121/1.4818773

50. Grose, J. H., & Mamo, S. K. (2010). Processing of temporal fine structure as a function of age. Ear and Hearing, 31, 755–760. 10.1097/AUD.0b013e3181e627e7

51. Guilleminot, P., & Reichenbach, T. (2022). Enhancement of speech-in-noise comprehension through vibrotactile stimulation at the syllabic rate. Proceedings of the National Academy of Sciences of the United States of America, 119(13), e2117000119. 10.1073/pnas.2117000119

52. He, D., Buder, E. H., & Bidelman, G. M. (2023). Effects of syllable rate on neuro-behavioral synchronization across modalities: Brain oscillations and speech productions. Neurobiology of Language, 4(2), 344–360. 10.1162/nol_a_00102

53. He, X., Raghavan, V. S., & Mesgarani, N. (2025). Neural speech tracking in noise reflects the opposing influence of SNR on intelligibility and attentional effort. Imaging Neuroscience, 3. 10.1162/IMAG.a.126

54. Heidari, A., Moossavi, A., Yadegari, F., Bakhshi, E., & Ahadi, M. (2018). Effects of age on speech-in-noise identification: Subjective ratings of hearing difficulties and encoding of fundamental frequency in older adults. Journal of Audiology & Otology, 22(3), 134–139. 10.7874/jao.2017.00304

55. Herrmann, B. (2025). Age-related differences in the impact of background noise on neural speech tracking. Neurobiology of Aging, 153, 10–20. 10.1016/j.neurobiolaging.2025.06.001

56. Hopkins, K., & Moore, B. C. J. (2009). The contribution of temporal fine structure to the intelligibility of speech in steady and modulated noise. The Journal of the Acoustical Society of America, 125(1), 442–446. 10.1121/1.3037233

57. Hopkins, K., & Moore, B. C. J. (2011). The effects of age and cochlear hearing loss on temporal fine structure sensitivity, frequency selectivity, and speech reception in noise. Journal of the Acoustical Society of America, 130, 334–349. 10.1121/1.3585848

58. Humes, L. E., Busey, T. A., Craig, J., & Kewley-Port, D. (2013). Are age-related changes in cognitive function driven by age-related changes in sensory processing? Attention, Perception, & Psychophysics, 75, 508–524. 10.3758/s13414-012-0406-9

59. Humes, L. E., Dubno, J. R., Gordon-Salant, S., Lister, J. J., Cacace, A. T., Cruickshanks, K. J., Gates, G. A., Wilson, R. H., & Wingfield, A. (2012). Central presbycusis: A review and evaluation of the evidence. Journal of the American Academy of Audiology, 23(8), 635–666. 10.3766/jaaa.23.8.5

60. Irsik, V. C., Johnsrude, I. S., & Herrmann, B. (2022). Age-related deficits in dip-listening evident for isolated sentences but not for spoken stories. Scientific Reports, 12(1), 5898. 10.1038/s41598-022-09805-6

61. Janssen, T., Steinhoff, H. J., & Böhnke, F. (1991). Zum entstehungsmechanismus der frequenzfolgepotentiale. Oto-Rhino-Laryngologia Nova, 1(1), 16–24. 10.1159/000312727

62. Kale, S., & Heinz, M. G. (2010). Envelope coding in auditory nerve fibers following noise-induced hearing loss. Journal of the Association for Research in Otolaryngology, 11(4), 657–673. 10.1007/s10162-010-0223-6

63. Keshavarzi, M., Kegler, M., Kadir, S., & Reichenbach, T. (2020). Transcranial alternating current stimulation in the theta band but not in the delta band modulates the comprehension of naturalistic speech in noise. Neuroimage, 210, 116557. 10.1016/j.neuroimage.2020.116557

64. Killion, M. C., Niquette, P. A., Gudmundsen, G. I., Revit, L. J., & Banerjee, S. (2004). Development of a quick speech-in-noise test for measuring signal-to-noise ratio loss in normal-hearing and hearing-impaired listeners. Journal of the Acoustical Society of America, 116(4 Pt 1), 2395–2405. 10.1121/1.1784440

65. Kochkin, S. (2010). MarkeTrak VIII: Consumer satisfaction with hearing aids is slowly increasing. The Hearing Journal, 63(1), 19–20,22,24,26,28,30–32. 10.1097/01.Hj.0000366912.40173.76

66. Lachaux, J. P., Rodriguez, E., Martinerie, J., & Varela, F. J. (1999). Measuring phase synchrony in brain signals. Human Brain Mapping, 8(4), 194–208. 10.1002/(SICI)1097-0193(1999)8:4<194::AID-HBM4>3.0.CO;2-C

67. Lai, J., Price, C. N., & Bidelman, G. M. (2022). Brainstem speech encoding is dynamically shaped online by fluctuations in cortical α state. Neuroimage, 263, 119627. 10.1016/j.neuroimage.2022.119627

68. Lentz, J. J., Humes, L. E., & Kidd, G. R. (2022). Differences in auditory perception between young and older adults when controlling for differences in hearing loss and cognition. Trends in Hearing, 26, 23312165211066180. 10.1177/23312165211066180

69. Macmillan, N. A., & Creelman, C. D. (2005). Detection theory: A user’s guide (2nd ed.). Lawrence Erlbaum Associates, Inc.

70. Mai, G., Tuomainen, J., & Howell, P. (2018). Relationship between speech-evoked neural responses and perception of speech in noise in older adults. The Journal of the Acoustical Society of America, 143(3), 1333–1345. 10.1121/1.5024340

71. Martin, J. S., & Jerger, J. F. (2005). Some effects of aging on central auditory processing. Journal of Rehabilitation Research and Development, 42(4 Suppl 2), 25–44. 10.1080/14992020500146450.

72. McClaskey, C. M. (2024). Neural hyperactivity and altered envelope encoding in the central auditory system: Changes with advanced age and hearing loss. Hearing Research, 442, 108945. 10.1016/j.heares.2023.108945

73. McClaskey, C. M., Dias, J. W., & Harris, K. C. (2019). Sustained envelope periodicity representations are associated with speech-in-noise performance in difficult listening conditions for younger and older adults. Journal of Neurophysiology, 122(4), 1685–1696. 10.1152/jn.00845.2018

74. Millman, R. E., Mattys, S. L., Gouws, A. D., & Prendergast, G. (2017). Magnified neural envelope coding predicts deficits in speech perception in noise. Journal of Neuroscience, 37(32), 7727–7736. 10.1523/jneurosci.2722-16.2017

75. Momtaz, S., Moncrieff, D., & Bidelman, G. M. (2021). Dichotic listening deficits in amblyaudia are characterized by aberrant neural oscillations in auditory cortex. Clinical Neurophysiology, 132(9), 2152–2162. 10.1016/j.clinph.2021.04.022

76. Moore, B. C. J., & Glasberg, B. R. (2001). Temporal modulation transfer functions obtained using sinusoidal carriers with normally hearing and hearing-impaired listeners. Journal of the Acoustical Society of America, 110(2), 1067–1073.

77. Moore, D. R., Edmondson-Jones, M., Dawes, P., Fortnum, H., McCormack, A., Pierzycki, R. H., & Munro, K. J. (2014). Relation between speech-in-noise threshold, hearing loss and cognition from 40–69 years of age. PloS One, 9(9), e107720. 10.1371/journal.pone.0107720

78. Nash, S. D., Cruickshanks, K. J., Klein, R., Klein, B. E. K., Nieto, F. J., Huang, G. H., Pankow, J. S., & Tweed, T. S. (2011). The prevalence of hearing impairment and associated risk factors: The Beaver Dam offspring study. Archives of Otolaryngology–Head & Neck Surgery, 137(5), 432–439. 10.1001/archoto.2011.15

79. Nimon, K., & Reio, T. G. (2011). Regression commonality analysis: A technique for quantitative theory building. Human Resource Development Review, 10(3), 329–340. 10.1177/1534484311411077

80. Nittrouer, S., & Boothroyd, A. (1990). Context effects in phoneme and word recognition by young children and older adults. Journal of the Acoustical Society of America, 87(6), 2705–2715. 10.1121/1.399061

81. Oostenveld, R., & Praamstra, P. (2001). The five percent electrode system for high-resolution EEG and ERP measurements. Clinical Neurophysiology, 112, 713–719. 10.1016/S1388-2457(00)00527-7

82. Parthasarathy, A., Datta, J., Torres, J. A., Hopkins, C., & Bartlett, E. L. (2014). Age-related changes in the relationship between auditory brainstem responses and envelope-following responses. Journal of the Association for Research in Otolaryngology, 15(4), 649–661. 10.1007/s10162-014-0460-1

83. Petersen, E. B., Wöstmann, M., Obleser, J., & Lunner, T. (2017). Neural tracking of attended versus ignored speech is differentially affected by hearing loss. Journal of Neurophysiology, 117(1), 18–27. 10.1152/jn.00527.2016

84. Pichora-Fuller, K. M., Alain, C., & Schneider, B. (2017). Older adults at the cocktail party. In J. C. Middlebrook, J. Z. Simon, A. N. Popper, & R. F. Fay (Eds.), Springer Handbook of Auditory Research: The Auditory System at the Cocktail Party (pp. 227–259). Springer-Verlag.

85. Picton, T. W., Alain, C., Woods, D. L., John, M. S., Scherg, M., Valdes-Sosa, P., Bosch-Bayard, J., & Trujillo, N. J. (1999). Intracerebral sources of human auditory-evoked potentials. Audiology & Neuro-otology, 4(2), 64–79. 10.1159/000013823

86. Picton, T. W., Woods, D. L., Baribaeu-Braun, J., & Healy, T. M. G. (1977). Evoked potential audiometry. Journal of Otolaryngology, 6(2), 90–119.

87. Presacco, A., Simon, J. Z., & Anderson, A. (2016a). Effect of informational content of noise on speech representation in the aging midbrain and cortex. Journal of Neurophysiology, 116(5), 2356–2367. 10.1152/jn.00373.2016

88. Presacco, A., Simon, J. Z., & Anderson, S. (2016b). Evidence of degraded representation of speech in noise, in the aging midbrain and cortex. Journal of Neurophysiology, 116(5), 2346–2355. 10.1152/jn.00372.2016

89. Price, C. N., & Bidelman, G. M. (2021). Attention reinforces human corticofugal system to aid speech perception in noise. Neuroimage, 235, 118014. 10.1016/j.neuroimage.2021.118014

90. Ratelle, D., & Tremblay, P. (2025). Neural tracking of continuous speech in adverse acoustic conditions among healthy adults with normal hearing and hearing loss: A systematic review. Hearing Research, 466, 109367. 10.1016/j.heares.2025.109367

91. Riecke, L., Formisano, E., Sorger, B., Başkent, D., & Gaudrain, E. (2018). Neural entrainment to speech modulates speech intelligibility. Current Biology, 28(2), 161–169.e165. 10.1016/j.cub.2017.11.033

92. Rogers, C. S., Jacoby, L. L., & Sommers, M. S. (2012). Frequent false hearing by older adults: The role of age differences in metacognition. Psychology and Aging, 27(1), 33–45. 10.1037/a0026231

93. Roque, L., Karawani, H., Gordon-Salant, S., & Anderson, S. (2019). Effects of age, cognition, and neural encoding on the perception of temporal speech cues. Frontiers in Neuroscience, 13. 10.3389/fnins.2019.00749

94. Salthouse, T. A., & Lichty, W. (1985). Tests of the neural noise hypothesis of age-related cognitive change. Journal of Gerontology, 40(4), 443–450. 10.1093/geronj/40.4.443

95. Schmitt, R., Meyer, M., & Giroud, N. (2022). Better speech-in-noise comprehension is associated with enhanced neural speech tracking in older adults with hearing impairment. Cortex, 151, 133–146. 10.1016/j.cortex.2022.02.017

96. Schneider, B. A., Daneman, M., & Pichora-Fuller, M. K. (2002). Listening in aging adults: From discourse comprehension to psychoacoustics. Canadian Journal of Experimental Psychology, 56(3), 139–152. 10.1037/h0087392

97. Sidiras, C., Iliadou, V. V., Nimatoudis, I., & Bamiou, D.-E. (2020). Absence of rhythm benefit on speech in noise recognition in children diagnosed with auditory processing disorder. Frontiers in Neuroscience, 14. 10.3389/fnins.2020.00418

98. Smith, M. L., Winn, M. B., & Fitzgerald, M. B. (2024). A large-scale study of the relationship between degree and type of hearing loss and recognition of speech in quiet and noise. Ear and Hearing, 45(4), 915–928. 10.1097/aud.0000000000001484

99. Strelcyk, O., & Dau, T. (2009). Relations between frequency selectivity, temporal fine-structure processing, and speech reception in impaired hearing. Journal of the Acoustical Society of America, 125(5), 3328–3345. 10.1121/1.3097469

100. Tarabichi, O., Kozin, E. D., Kanumuri, V. V., Barber, S., Ghosh, S., Sitek, K. R., Reinshagen, K., Herrmann, B., Remenschneider, A. K., & Lee, D. J. (2018). Diffusion tensor imaging of central auditory pathways in patients with sensorineural hearing loss: A systematic review. Otolaryngology–Head and Neck Surgery, 158(3), 432–442. 10.1177/0194599817739838

101. te Rietmolen, N., Strijkers, K., & Morillon, B. (2025). Moving rhythmically can facilitate naturalistic speech perception in a noisy environment. Proceedings of the Royal Society B: Biological Sciences, 292(2044), 20250354. 10.1098/rspb.2025.0354

102. van Bree, S., Sohoglu, E., Davis, M. H., & Zoefel, B. (2021). Sustained neural rhythms reveal endogenous oscillations supporting speech perception. PLoS Biol, 19(2), e3001142. 10.1371/journal.pbio.3001142

103. Vanthornhout, J., Decruy, L., Wouters, J., Simon, J. Z., & Francart, T. (2018). Speech intelligibility predicted from neural entrainment of the speech envelope. Journal of the Association for Research in Otolaryngology, 19, 181–191. 10.1007/s10162-018-0654-z

104. Voytek, B., & Knight, R. T. (2015). Dynamic network communication as a unifying neural basis for cognition, development, aging, and disease. Biological Psychiatry, 77(12), 1089–1097. 10.1016/j.biopsych.2015.04.016

105. Voytek, B., Kramer, M. A., Case, J., Lepage, K. Q., Tempesta, Z. R., Knight, R. T., & Gazzaley, A. (2015). Age-related changes in 1/f neural electrophysiological noise. The Journal of Neuroscience, 35(38), 13257–13265. 10.1523/jneurosci.2332-14.2015

106. Wallaert, N., Moore, B. C., Ewert, S. D., & Lorenzi, C. (2017). Sensorineural hearing loss enhances auditory sensitivity and temporal integration for amplitude modulation. Journal of the Acoustical Society of America, 141(2), 971. 10.1121/1.4976080

107. Wilsch, A., Neuling, T., Obleser, J., & Herrmann, C. S. (2018). Transcranial alternating current stimulation with speech envelopes modulates speech comprehension. Neuroimage, 172, 766–774. 10.1016/j.neuroimage.2018.01.038

108. Wingfield, A., & Tun, P. A. (2001). Spoken language comprehension in older adults: Interactions between sensory and cognitive change in normal aging. Seminars in Hearing, 22(03), 287–302. 10.1055/s-2001-15632

109. Wingfield, A., Tun, P. A., & McCoy, S. (2005). Hearing loss in older adulthood: What it is and how it interacts with cognitive performance. Current Directions in Psychological Science, 14(3), 144–148. 10.1111/j.0963-7214.2005.0035

110. Wong, P. C. M., Jin, J. X., Gunasekera, G. M., Abel, R., Lee, E. R., & Dhar, S. (2009). Aging and cortical mechanisms of speech perception in noise. Neuropsychologia, 47, 693–703. 10.1016/j.neuropsychologia.2008.11.032

111. Zobel, B. H., Wagner, A., Sanders, L. D., & Başkent, D. (2019). Spatial release from informational masking declines with age: Evidence from a detection task in a virtual separation paradigm. The Journal of the Acoustical Society of America, 146(1), 548–566. 10.1121/1.5118240

